# Production of Insoluble Starch-Like Granules in *Escherichia coli* by Modification of the Glycogen Synthesis Pathway

**DOI:** 10.1101/841023

**Authors:** Joseph J. White, Natasha Cain, Christopher E. French

**Author notes:** Address correspondence to Joseph J. White,. Department of Molecular Evolution, Centro de Astrobiología (CSIC-INTA), 28850 Torrejón de Ardoz (Madrid).

## Abstract

While investigating the conversion of cellulosic biomass to starch-like materials for industrial use, it was observed that the overexpression of native ADP-glucose pyrophosphorylase GlgC in *Escherichia coli* led to the formation of insoluble polysaccharide granules within the cytoplasm, occupying a large fraction of the cell volume, as well as causing an overall increase in cellular polysaccharide content. TEM microscopy revealed that the granules did not have the lamellar structure of starch, but rather an irregular, clustered structure. On starvation, cells overexpressing GlgC appeared unable to fully degrade their polysaccharide material and granules were still clearly visible in cultures after 8 days of starvation. Interestingly, the additional overexpression of the branching enzyme GlgB eliminated the production of granules and led to a further increase in cellular polysaccharides. GlgC is generally thought to be responsible for the rate-limiting step of glycogen synthesis. Our interpretation of these results is that excess GlgC activity may cause the elongation of glycogen chains to outpace the addition of side branches, allowing the chains of adjacent glycogen molecules to reach lengths at which they spontaneously intertwine, forming dense clusters that are largely inaccessible to the host. However, upon additional upregulation of the GlgB branching enzyme, the branching of the polysaccharide is able to keep speed with the synthesis of linear chains, eliminating the granule phenotype. This study suggests potential avenues for increasing bacterial polysaccharide production and recovery.

**Importance:** In this work, the polysaccharide stores of *Escherichia coli* were altered through the addition of extra copies of the bacteria’s own polysaccharide synthesis genes. In this way, bacteria were created that produced over twice the level of storage polysaccharide as a control strain, in the form of a granule that could potentially facilitate easy harvest. Another form of mutant *Escherichia coli* was created that produced over seven times the normal level of storage polysaccharide, and also grew to higher cell densities in liquid culture. In addition to increasing our understanding of glycogen synthesis, it is proposed that similarly modified bacteria, grown on inexpensive waste materials, may be a useful source of starch-like polysaccharides for industrial or agricultural use. In particular, the use of cyanobacterial glycogen as a carbon source for biofuels has recently been gaining interest, and the work presented here may well be applicable in this field.

## Introduction

Starch granules accumulated by plant cells form the basis of a large part of the human food supply, and are a valuable resource in many industries, including paper and biofuels. Recent years have seen enormous volumes of grain diverted to bioethanol production; reportedly, in 2013-2014, 40% of the US maize crop was used for biofuel production (1). According to some reports, this was a factor in the food price riots in many parts of the world in 2008 (2, 3). Furthermore, it has been suggested that the land clearance necessary for growing crops used as biofuel may produce more CO_2_ emissions than are saved through the reduction of fossil fuel use (4, 5).

Whether or not all this is the case, it is clear that as the human population increases, it will be increasingly unacceptable to divert human-grade food materials to non-food uses. Thus, a non-food based source of starch-like polymers is required. One current plan to reduce diversion of food materials would appear to be a gradual change to ‘second generation’ biofuels, such as ethanol generated directly from abundant cellulosic waste material by genetically modified strains of *Saccharomyces cerevisiae*, *Escherichia coli*, or other organisms such as *Geobacillus thermoglucosidasius* (6–11). Despite the expenditure of vast resources, and significant government subsidies and other incentives, cellulosic ethanol production currently remains at a rather small scale. One apparently unexplored alternative would be conversion of sugars from cellulosic material to an easily degradable starch-like form, which could be incorporated into the current, very large scale grain-based ethanol production system. Such a two-stage process might have significant advantages in terms of process kinetics, since conversion of cellulosic biomass to sugars is unavoidably a ‘slow’ step compared to fermentation of sugars to ethanol. Initial slow conversion of cellulosic polymers to an insoluble but easily degradable starch-like material, which can be easily recovered from the fermentation medium, washed, transported, and stored for later use, followed by rapid saccharification and fermentation using standard, highly efficient processes, may alter the economics of cellulosic ethanol production. In addition, microbial starch-like polymers may also be useful in other industries, and might even be able to partially substitute for the use of grain in feeding livestock, thus freeing up more grain for human food use.

For these reasons the production of insoluble starch-like polymers has been investigated in bacteria. In general the production of true starch is limited to plants and green and red alga, though it has now also been identified in at least one strain of subgroup V diazotrophic cyanobacteria (12, 13). Common bacteria such as *Escherichia coli* produce glycogen, which has a similar structure to the amylopectin fraction of starch, that is to say α-1,4-linked glucose chains with α-1,6-linked side branches, but with a higher branching level of around 9%. The branching level of amylopectin is below 6% and occurs discontinuously, in tiers, thereby creating alternating amorphous and crystalline lamella (14, 15). The crystalline lamella are likely composed of unbranched glucan chains around 12 to 20 glucosyl residues in length, that spontaneously pair together forming double helices with their neighbours, while the amorphous lamella are thought to be the recurrent regions of the molecule that contain the branch points, and as such are prevented from forming helices by the interference of the α-1,6-bonds. It is further supposed that the double-helices of the crystalline lamella twist into superhelices, although the details of this have yet to be elucidated (16–18).

The amylose fraction generally makes up around 15% of the starch crystal in the Chloroplastida and is thought to be interspersed within the amorphous lamella of the amylopectin, although its exact location is still under debate (19, 20). It is far simpler than amylopectin, being mostly unbranched chains of α-1,4 linked glucosyl residues. It is also far smaller, with molecular weight estimates varying between 10^5^ and 10^6^ Daltons (18). Because of its sparsity of branches, amylose twists itself into single helices, with each coil of the helix thought to be 6 glucosyl residues long (21).

Iodine is often used in the colourimetric assaying of starch, and it is the single helices of the amylose fraction that react to form the intense blue-black colour typical of this assay. The resulting colour depends on the length of the helix-iodine complex, going from brown to red to blue with increasing length (20, 22, 23). Amylopectin also stains with iodine solution, though with a much lower affinity (<1%, compared to around 20% with amylose) since iodine does not complex with the double-helices of the crystalline lamella. The amylopectin-iodine complex therefore tends to be very pale, with a reddish-purple colour. Glycogen, meanwhile, generally forms a deep red complex with iodine, suggesting an abundance of shorter single helices, although this varies considerably among different sources due to glycogen’s lower structural organisation (24).

Due to the high branching level of glycogen, its granules are thought to be limited in size to around 42 nm diameter by steric considerations, and are highly soluble, whereas starch granules, with their tiered growth, may be hundreds of micrometres in diameter and are insoluble, providing a denser energy store (25, 26). The main chains of glycogen are synthesised by glycogen synthase (GlgA), which adds sugar residues in the form of ADP-glucose, produced from glucose-1-phosphate and ATP by ADP-glucose pyrophosphorylase (GlgC). The activity of GlgC is therefore generally considered to be the first committed step and also the rate-limiting step in glycogen synthesis (14). Meanwhile, the α-1,6-linked side chains are added by branching enzyme GlgB, which cleaves about four residues of a chain off the terminal end and attaches them to the 6-hydroxyl of a residue further down the chain. The highly branched structure of glycogen allows optimum accessibility of glucose residues to the degradative enzymes glycogen phosphorylase (GlgP) and glycogen debranching enzyme (GlgX) (27) (figure 1).

**Figure 1:**
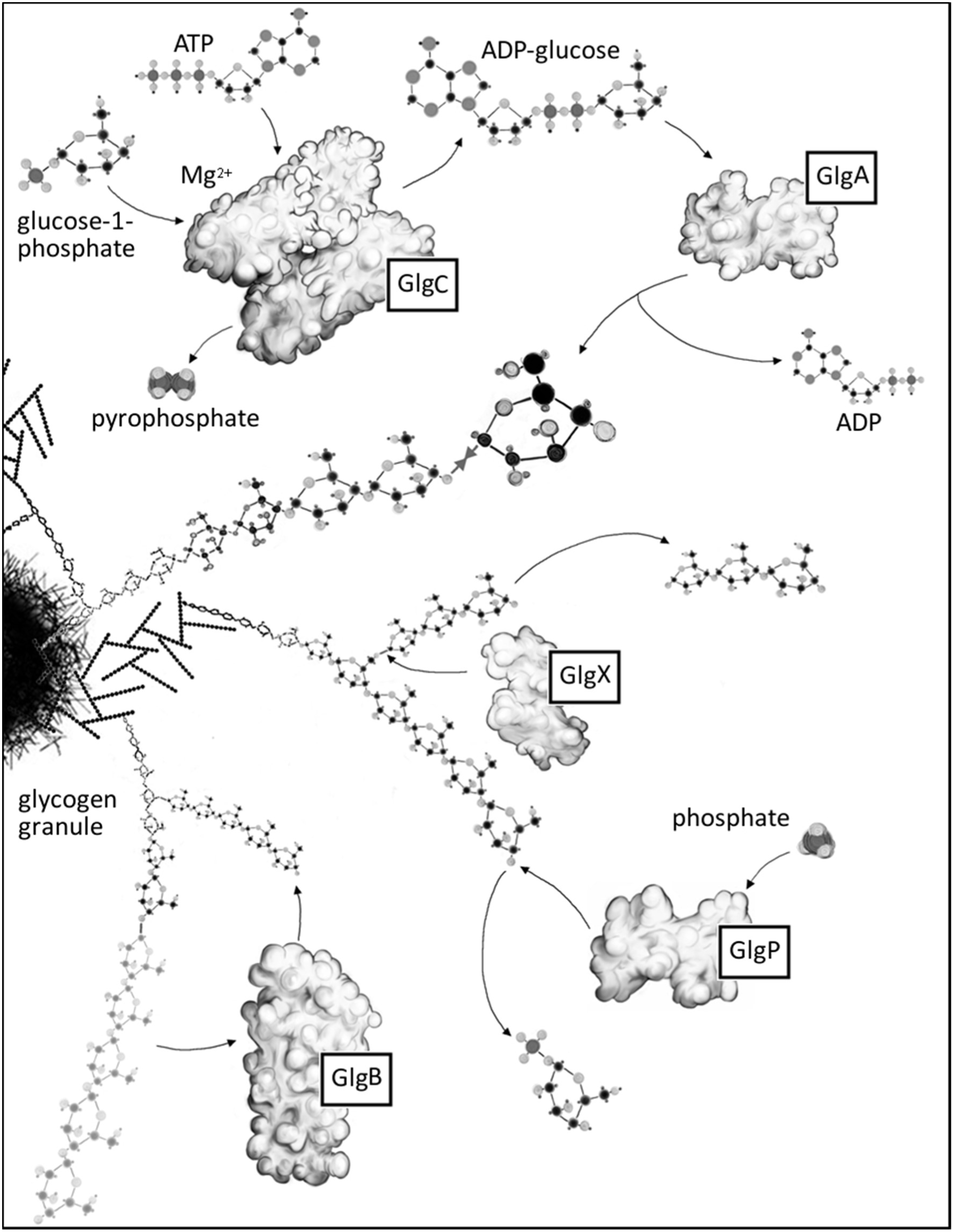
**Glycogen metabolism in *E. coli*.** GlgC and GlgA work in tandem to grow the non-reducing ends of external glucan chains one glucose unit at a time, while GlgP removes glucose residues from the non-reducing ends of the chains, until its activity is cut short by proximity to the branches created by GlgB. GlgX can sever the α-1,6 bonds at the fork of these branches, but only on branches shorter than those GlgB has originally attached, thereby preventing the enzymes from falling into a futile cycle of successive branching and debranching (Cenci et al., 2014).

To investigate the possibility of converting bacterial glycogen to an insoluble starch-like form for easy recovery and processing, this research initially sought to upregulate glycogen production. In line with the idea that GlgC catalyses the rate-limiting step of glycogen synthesis, it is known that the upregulation of glycogen production can be accomplished by the overexpression of GlgC, or more particularly, a mutant form, GlgC16 (G335D), which is resistant to normal feedback inhibition by AMP (28). However, on overexpression of GlgC in *E. coli* MG1655 JM109, we observed large insoluble bodies within the cells, apparently composed of glucose polymers. To the best of our knowledge, this phenomenon has not previously been reported, although glycogen synthesis has been extensively studied. Here the preliminary characterization of these inclusions is described.

## Results

In order to create cells with a higher polysaccharide content, GlgC and GlgC16 were overexpressed in *Escherichia coli* JM109 from a high copy number plasmid, pSB1C3, using a *lac* promoter (pJW-glgC and pJW-glgC16, respectively). The same plasmid with *lac* promoter and *lacZ*’ alone (pJW-lacZ) was used as a control (table 3). Cells bearing pJW-glgC or pJW-glgC16 grown in the presence of lactose were found to contain higher levels of polysaccharide than controls as expected and, correspondingly, to stain a darker colour with iodine (figure 2). Cells were treated with CuSO_4_ and H_2_O_2_ in order to stabilise the colour change with iodine (29). The effect of this treatment on the intensity of staining of starch-iodine reactions was investigated to ensure it did not interrupt the linear trend of colour intensity to polysaccharide content, and was found not to interrupt this linearity.

**Figure 2:**
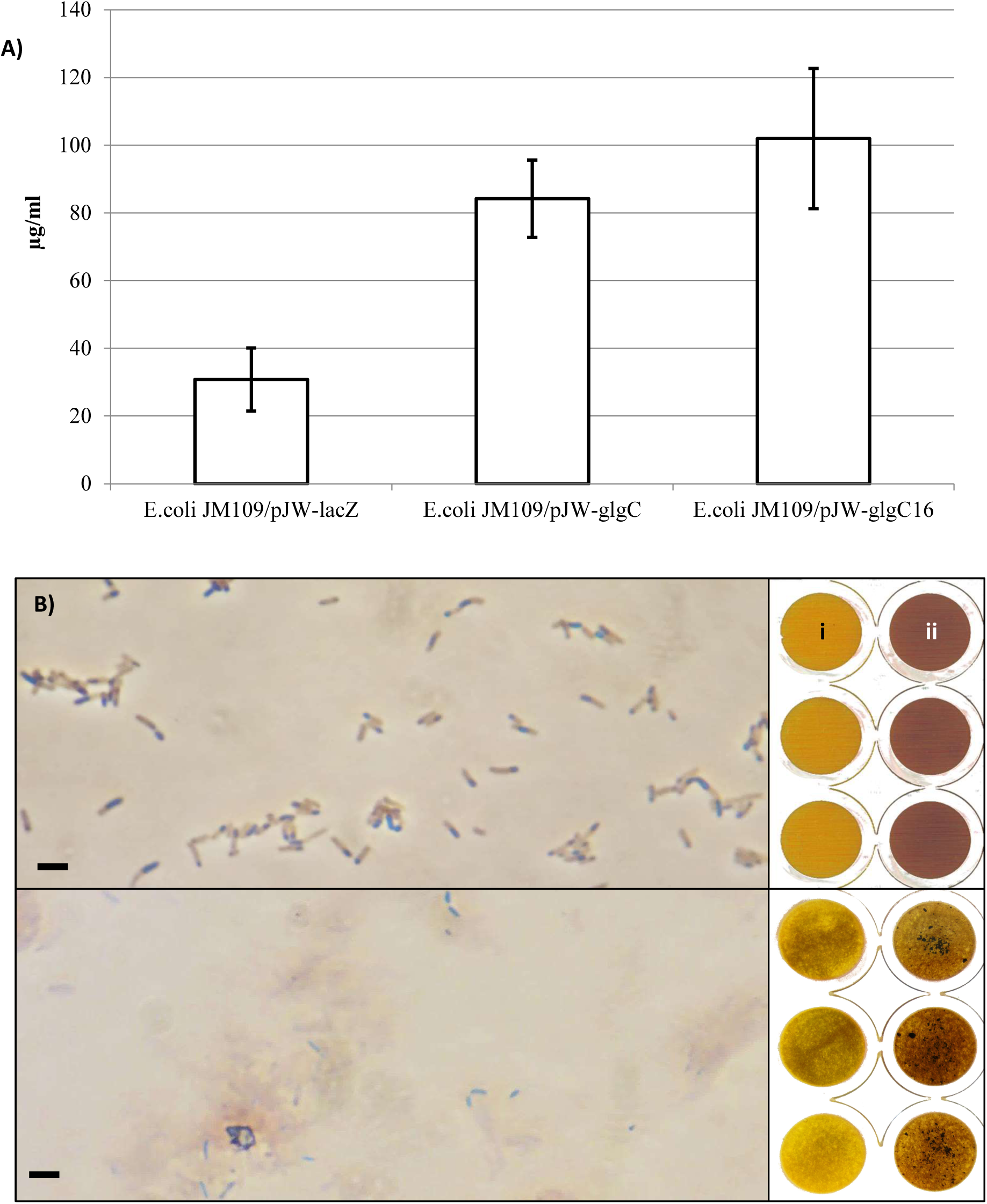
**(A) total hexose sugar content of *E. coli* JM109/pJW-lacZ versus *E. coli* JM109/pJW-glgC and *E. coli* JM109/pJW-glgC16, f**rom anthrone assays performed on cultures grown in a manner expected to maximise their polysaccharide content. Error bars represent the standard error of the mean when n=3. **(B) Phase contrast light microscopy showing ‘inclusion body’ phenotype in iodine stained *E. coli* JM109/pJW-glgC16 cells before (top) and after (bottom) lysis with KOH, with iodine assay results of the same cultures, pre and post lysis with KOH.** Cultures were grown overnight in LB supplemented with IPTG and 1% lactose. Lysis was achieved with 30% KOH and 5 minutes boiling. Scale bar represents 5 µm. Column ‘i’: *E. coli* JM109/pJW-lacZ. Column ‘ii’: *E. coli* JM109/pJW-glgC.

Unexpectedly, cells transformed with either pJW-glgC or pJW-glgC16 were also found to contain large inclusions, clearly visible under phase contrast microscopy, which were absent in cells containing a control plasmid. These inclusions stained darkly with iodine against the unstained cell body, appearing vivid blue under phase contrast (figure 2B) or brown under light-field microscopy (not shown). The iodine-staining material was also seen to be present in the cell debris rather than the soluble fraction (figure 2B). SDS-PAGE analysis showed no large new bands in the cytoplasm or cell debris compared to controls, indicating that the inclusions were not composed of misfolded protein.

It was hypothesised that an increased supply of ADP-glucose provided by the upregulation of *glgC* allowed glycogen synthase activity to reach a previously unrealised potential and outstrip the activity of the branching enzyme. To test this hypothesis pJW-glgCB was constructed, bearing *glgC* followed by *glgB*, both under the control of the *lac* promoter (table 3). In line with the prediction, *E. coli* JM109/pJW-glgCB produced increased levels of polysaccharide compared to the control, although unexpectedly the increase was found to be far greater still than for those cells transformed with the additional *glgC* alone (figure 3A). Also in line with predictions, cells transformed with pJW-glgCB were seen to stain the typical red-brown of glycogen in reaction with iodine (figure 3, B and C) and did not contain visible inclusions by light microscopy. All cultures were analysed by transmission electron microscopy, where both of the novel tranformants showed a marked phenotype different from controls (figure 4). The inclusions found in *E. coli* JM109/pJW-glgC cells were seen to have a granular internal structure. Cells transformed with pJW-glgCB were seen to contain an abundance of inclusive matter dispersed throughout the cells. In the case of both transformants, the inclusive matter was observed in around a third of cells.

**Figure 3:**
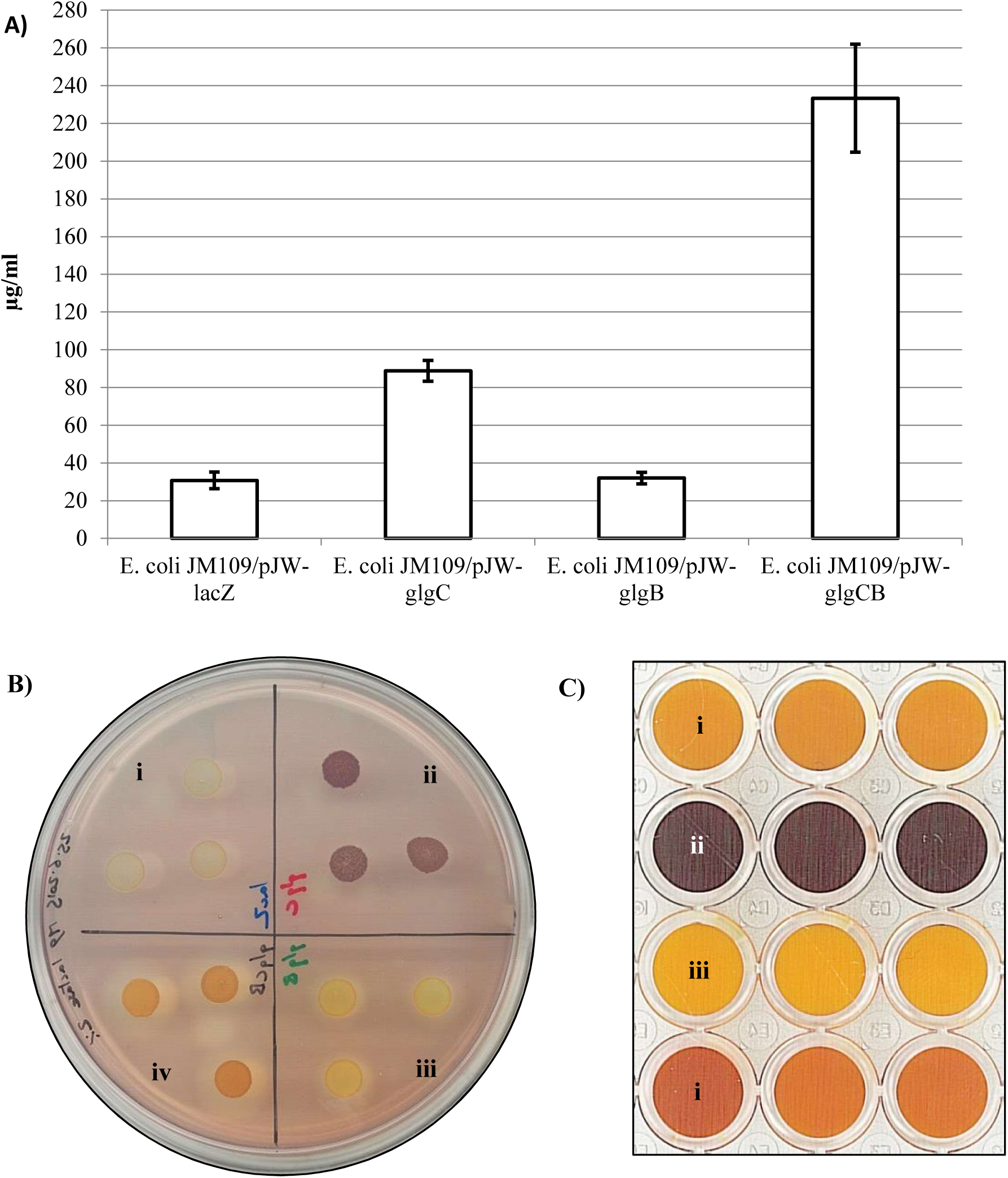
**(A) Total hexose sugar content of transformed JM109 cells.** Anthrone assays performed on cultures equalised to OD_λ600nm_ 3.5, after around 30 hours growth in Kornberg medium followed by M9, in a manner expected to maximise their polysaccharide content. Error bars represent the standard error of the mean when n=3. **(B & C) Iodine assays of transformed JM109.** B) patches from cultures grown to stationary phase in LB supplemented with lactose & IPTG, spotted onto an M9 plate, grown overnight at 37°C and flooded with Lugol’s iodine (0.2%).; C) Cell pellets from 1.4 ml of the same cultures used in image B, pelleted and resuspended in 200 µl PBS, plus 50 µl CuSO_4_ (100 mM), 50 µl H_2_O_2_ (6%) and 25 µl Lugol’s iodine (0.2%), transferred to the wells of a 48-well plate and imaged using a flatbed scanner. i) *E. coli* JM109/pJW-lacZ; ii) *E. coli* JM109/pJW-glgC; iii) *E. coli* JM109/pJW-glgB; iv) *E. coli* JM109/pJW-glgCB.

**Figure 4:**
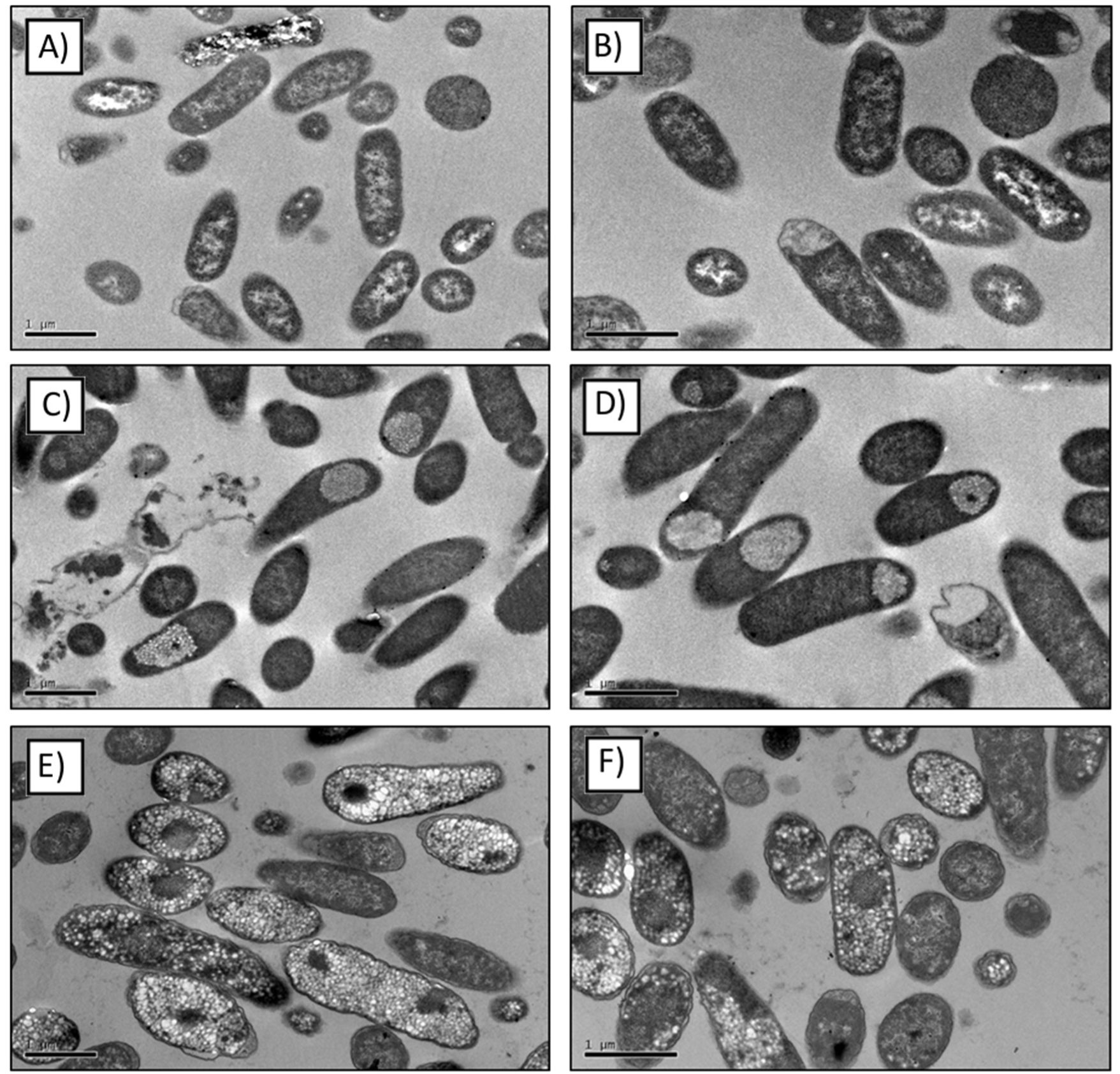
**Transmission electron micrographs (TEMs) of *E. coli* strain JM109 transformed with the pJW-lacZ ‘control’ plasmid (A,B), pJW-glgC16 plasmid (C,D) or pJW-glgCB plasmid (E,F).** Many of the ‘control’ cells show negatively-stained inclusive matter. However, it is considered that the defined granular bodies visible within many of the pJW-glgC16 and pJW-glgCB transformed cells are distinct, both from the control and from each other, and in the case of pJW-glgC16, correspond to those bodies found to stain with iodine when viewed under light microscopy. Images are selected from 14 pJW-lacZ, 20 pJW-glgC16 and 25 pJW-glgCB transmission electron micrographs captured, and are considered to be representative.

It was also observed that *E.coli* JM109/pJW-glgCB cultures reached higher densities than those of the control. To investigate this further, growth curves were obtained for the different transformants in both LB and modified Kornberg medium, substituted with lactose and IPTG (figure 5). *E. coli* JM109/pJW-glgCB showed slower initial growth than the control for the first five hours growth in LB, but then continued to increase in cell density and plateaued at a higher density than any of the other transformants. Cells transformed with pJW-glgC showed an even slower initial growth rate, only reaching the same density as the control after around eight hours growth in LB, and were never seen to exceed the cell density of the control in this medium. A similar pattern was seen in the modified Kornberg medium, although all cultures reached a higher final density and the differences between final culture densities were less clear. Cells transformed with pJW-glgB showed an almost identical growth curve to the control in all circumstances.

**Figure 5:**
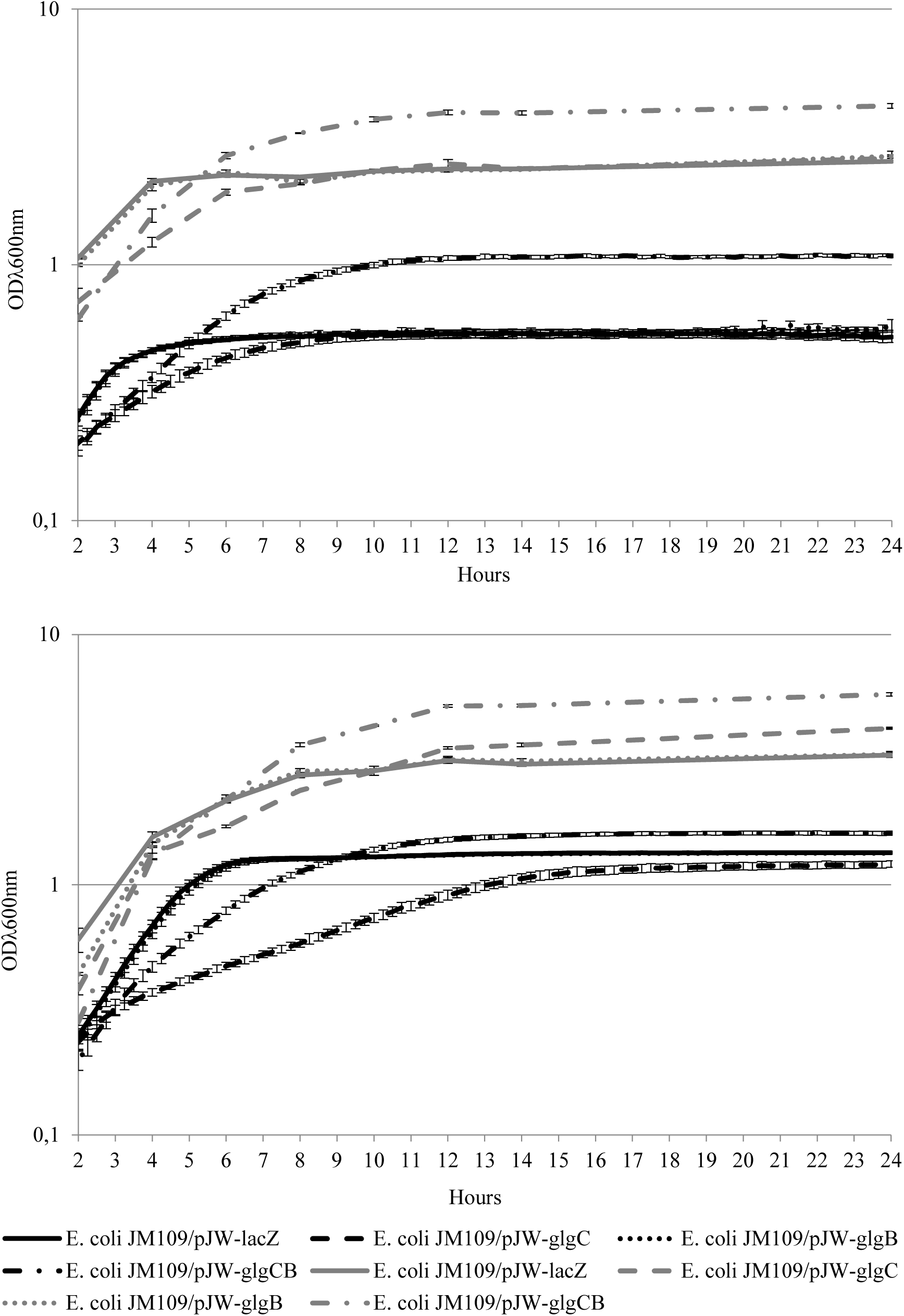
**Growth curves for the four transformants, grown in LB media (A) or modified Kornberg media (B), supplemented with lactose and IPTG, for 24 hours.** Grey lines show 50 ml shake flask readings. Error bars represent the standard error of the mean when n=3. Black lines show 96-well plate readings from 3 plates. Error bars represent the standard error of the mean when n=6.

It was subsequently tested whether the polysaccharides were accessible to cells under starvation conditions. Cultures of all transformants were grown to the same optical density in a way that was expected to maximise their polysaccharide content, before the cells were washed and resuspended in minimal medium without added carbon source. Culture density readings were taken at intervals over 216 hours, while anthrone assays were performed on aliquots of the cultures at the same time points to determine how much of the polysaccharide present in the original cell pellets was metabolized over time (figure 6, A and B). All cultures showed a decrease in culture density and polysaccharide content. In line with expectations, *E. coli* JM109/pJW-glgC showed a much greater reduction in culture density than the control over this time, and was never observed to fully deplete their polysaccharide stores. However, the greatest reduction in culture density was seen in *E. coli* JM109/pJW-glgCB. Furthermore, these cultures were also never able to fully deplete their polysaccharide stores, even when the experiment was repeated over 264 hours to clarify ambiguous results (figure 6C). In all cases *E. coli* JM109/pJW-glgB was once again observed to behave in a way similar to the control. Protein assays were performed on all cultures for the final two time points of the starvation experiment and were seen to corroborate the optical density readings. Cultures were also observed under light microscopy after staining with iodine at each time point of the starvation experiment. Inclusion bodies were observed at a similar ratio within the *E. coli* JM109/pJW-glgC cells throughout the course of both experiments (Figure 7).

**Figure 6:**
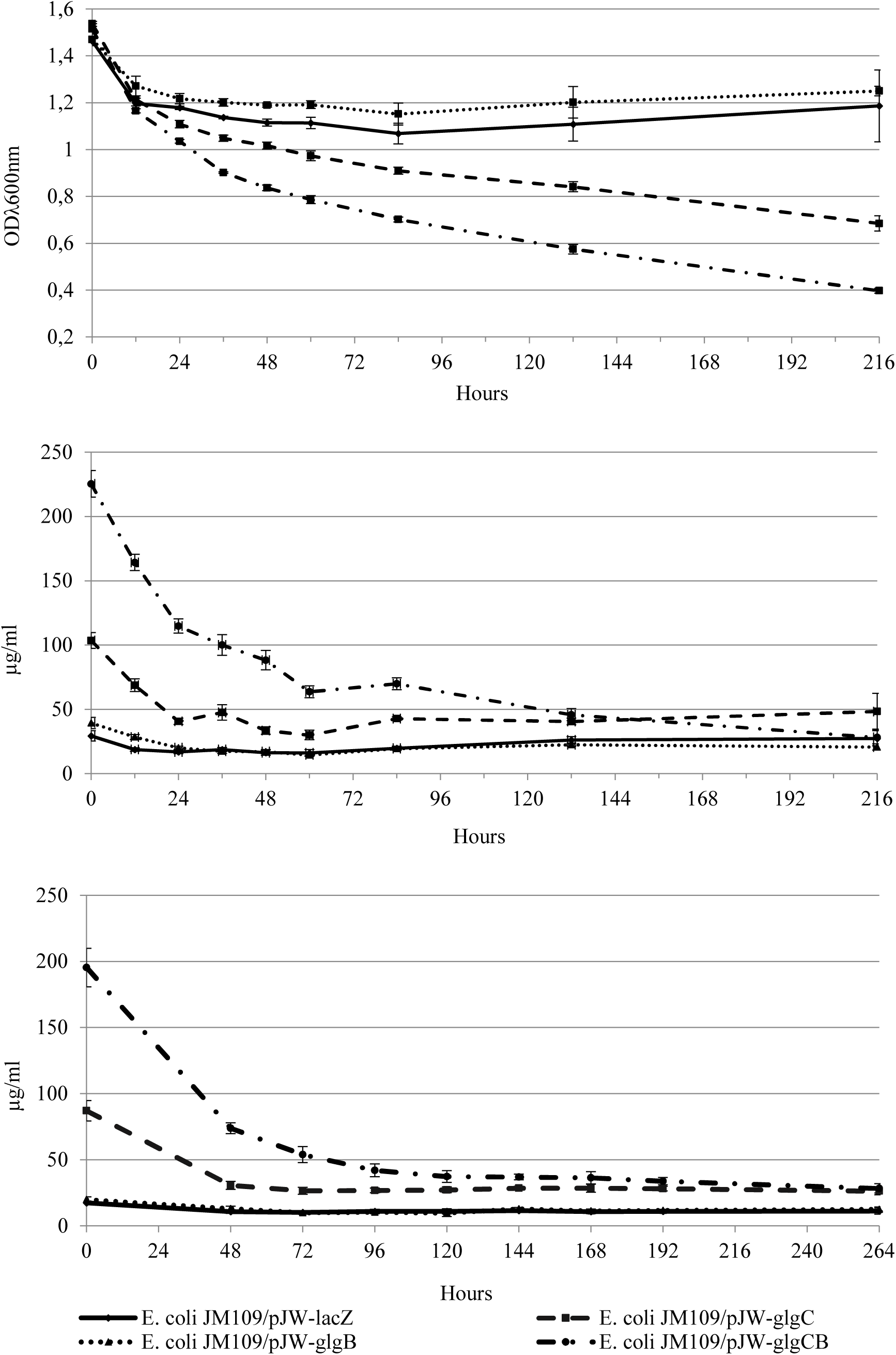
**(A) Growth curves for transformed JM109 over 216 hours of starvation conditions**. Starting from cell densities of 1.5 OD_λ600nm_. **(B) Total hexose sugar content of cultures over the course of the starvation experiment. (C) Repeat of total hexose sugar content experiment.** Error bars represent the standard error of the mean when n=3.

**Figure 7:**
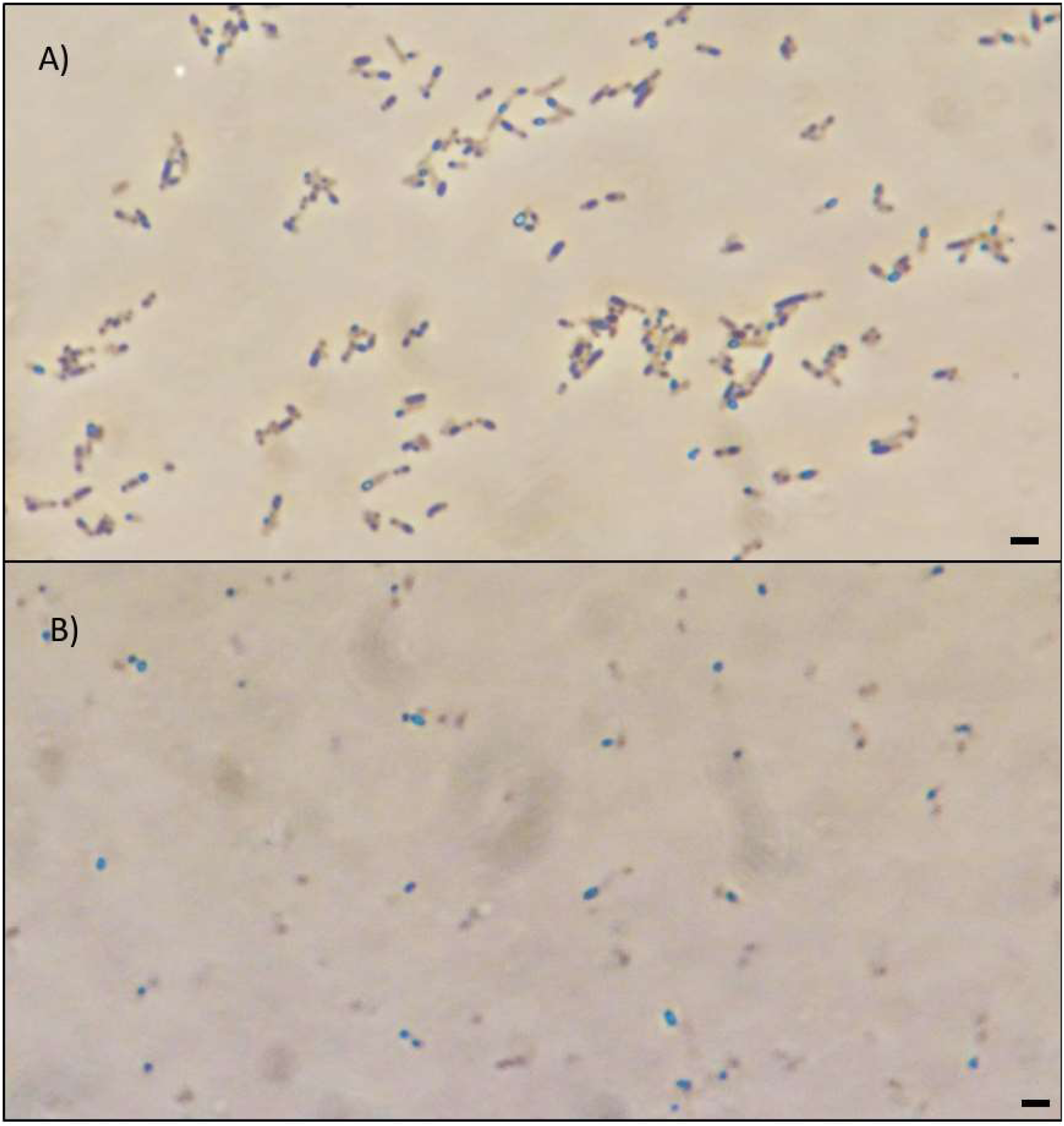
**Light microscopy of iodine stained, *E. coli* JM109/pJW-glgC before and after starvation.** From cultures of *E. coli* JM109/pJW-glgC used in the starvation experiment, imaged on day 1 (A) and day 8 (B), showing clear inclusion bodies in both cases. Scale bar represents 5 µm (approx.)

## Discussion

This work showed that the transformation of *E. coli* MG1655 JM109 with additional copies of its native *glgC*, expressed on a high copy number plasmid under a *lac* promoter, led to the formation of inclusion bodies within the cells when they were grown in culture supplemented with lactose and IPTG. Inclusion bodies were also observed in the same transformants when grown in media supplemented with glucose in place of lactose, although to a lesser extent, which is thought to be due to the inhibitory effect of glucose on the *lac* promoter (data not shown). The same ‘inclusion body’ phenotype was not observed in cells transformed with the same plasmid but lacking *glgC*, under any of the growth conditions tested. When present, inclusion bodies were observed to stain a deep brown in reaction with iodine, suggestive of an abundance of single-helices shorter than those found in glycogen, which tends to stain red. SDS-PAGE analysis of the *E. coli* JM109/pJW-glgC16 cells suggested that the inclusions observed were not composed of a single over-expressed protein.

Anthrone assays performed on transformants in this work have been consistent in showing a higher hexose sugar content for *glgC16* and *glgC* transformed cells than for a control. It was hypothesised that, since GlgC activity normally performs the rate-limiting step of glycogen synthesis, upregulating its expression in JM109 *E. coli* grown in sugar-rich media leads to the appearance of inclusion bodies in many of the cells, as well as a higher average intracellular concentration of sugar than in a control group grown under the same conditions, because GlgC increases the substrate availability for GlgA, the glycogen synthase. This increased substrate is thought to allow the glycogen synthase to meet a previously unrealised potential and synthesise the linear glucan chains within the polysaccharide granule at a faster rate than GlgB can add branches to them, so that they grow long unbranched chains able to spontaneously form double-helices, and that this interweaving of adjacent chains is causing the aggregation of granules into the large inclusion bodies observed. These double-helices would not bind iodine. However, since overall polysaccharide content is also increased approximately twofold by this manipulation, we suggest that it also leads to the appearance of abundant short single helices, giving the deep brown stain observed. The granular internal structure observed in the inclusions under TEM, which are unlike the lamellar structure seen in starch granules, also supports the hypothesis that they are formed of clustered glycogen-like polysaccharides.

If this hypothesis were correct, it was thought that the addition of *glgB* to the same plasmid as *glgC* would allow the branching of the polysaccharide to keep speed with the synthesis of linear chains. This was expected to stop the formation of inclusion bodies, since unbranched regions of glucan chains would no longer be able to grow long enough to wind together into double-helices, and furthermore the steric interference caused by the dense packing of glucan branches, which theoretically limits the growth of native glycogen granules, would be reintroduced. However, the cells were still expected to show the high sugar content seen in those transformed with *glgC*, since the glycogen synthase would still be able to synthesise chains at the increased rate. All these phenotypes were observed, with the added observation that, rather than synthesising the same high levels of polysaccharide as the *glgC* transformed cells, those transformed with both *glgC* and *glgB* showed more than double their total sugar content.

Furthermore, this work has shown that *E. coli* JM109/pJW-glgCB grow to a significantly higher culture density than *E. coli* JM109/pJW-lacZ grown under the same conditions. However, contrary to predictions, they have also proved to be far more vulnerable when exposed to starvation conditions, with culture densities dropping to less than one third of the control after nine days (the optical density readings that led to this conclusion correlate well with protein assays from the same cultures, suggesting they give an accurate measure of biomass rather than, for example, being an artefact of increased light scattering caused by the inclusion bodies). The excess storage sugar is therefore perhaps an added stress to these cells under such conditions. *E. coli* JM109/pJW-glgC were also more vulnerable to starvation conditions than the *E. coli* JM109/pJW-lacZ control, and also showed the slowest growth rate of any of the transformants. If indeed the inclusion bodies accumulated during prior growth in carbon-rich media are formed of polysaccharides, it seems as though these transformants were incapable of digesting them. The presence of inclusion bodies within *E. coli* JM109/pJW-glgC transformants at the beginning and end of the starvation period is felt to support this hypothesis. Although further testing is needed, these findings could have implications for the synthesis of polysaccharides for industry, since large inclusion bodies that the bacteria are unable to digest should be easier to harvest than native glycogen granules.

Starvation experiments additionally seemed to suggest that *E. coli* JM109/pJW-glgCB contained hexose sugars it was unable to digest, since anthrone assays for this transformant showed its total hexose sugar content level off at around the same concentration as that of the *E. coli* JM109/pJW-glgC. No inclusion bodies are observed in these cells under the microscope, although the cells show a dense concentration of inclusive matter under TEM. This is presumed to be glycogen. It is not clear why these transformants are unable to fully digest their hexose sugar content.

*E. coli* lack the dikinases (Glucan Water Dikinase and Phosphoglucan Water Dikinase) that are thought to aid in starch degradation by starch metabolising organisms, possibly by prising the α-1,4 linked glucan chains out of their double-helices in order to make them accessible to water soluble enzymes such as phosphorylase and debranching enzyme (GlgP and GlgX, respectively, in *E. coli*) (30, 31). Therefore the addition of *gwd* and *pwd* to the battery of transgenes expressed in these cells may allow for the digestion of the inclusion bodies. If so, this would support the theory that the additional *glgC* is leading to the growth of longer unbranched regions of glucan chains, which then spontaneously twist into double-helices with adjacent chains, causing an aggregation of polysaccharide.

In summary, this work demonstrates that simple modifications to the native glycogen synthesis machinery in *E. coli* can lead to the production of insoluble polysaccharide granules which appear indigestible under starvation conditions. Such polysaccharides may be easily recovered from culture media, and as such might represent a useful industrial method for capturing sugars released from cellulosic materials, in a form which can easily be converted to glucose for subsequent processing to generate biofuels or chemical feedstocks. Such ‘pseudo-starch’, if generated from non-digestible biomass or waste sugar materials, may also be tested as feed supplement for livestock, potentially releasing large amounts of grain for human food use.

## Materials and Methods

### Chemical transformation

*E. coli* JM109 competent cells were prepared and transformed as described by Chung et al. (32).

### Growth to maximise polysaccharide content

Overnight cultures (grown in LB (table 1), 37 °C, 200 rpm) were diluted 1:100 and grown to exponential growth phase (0.5 –0.6 OD_λ600nm_) in modified Kornberg Medium (table 1) with appropriate selection at 37°C and 200 rpm. Cultures were then supplemented with Isopropyl β-D-1-thiogalactopyranoside (IPTG) to 1 mM in order to induce P_lac_, and incubated for a further 20 hours at 22 °C and 200 rpm. They were then centrifuged at 3000 ×g for 15 minutes at room temperature, the supernatant was discarded and the pellet washed with M9 ×1 solution (table 1). Cell pellets were then resuspended in M9 medium with IPTG (90 mg/l), chloramphenicol (40 mg/l) and lactose (20 g/l) and incubated for a further 4 hours at 37 °C and 200 rpm (modified from Sundberg et al., 33). Cultures could then be spun down at 3000 ×g for 15 minutes at room temperature, washed twice in PBS (table 1) and equalised for optical density at 600nm, prior to analysis.

**Table 1:**
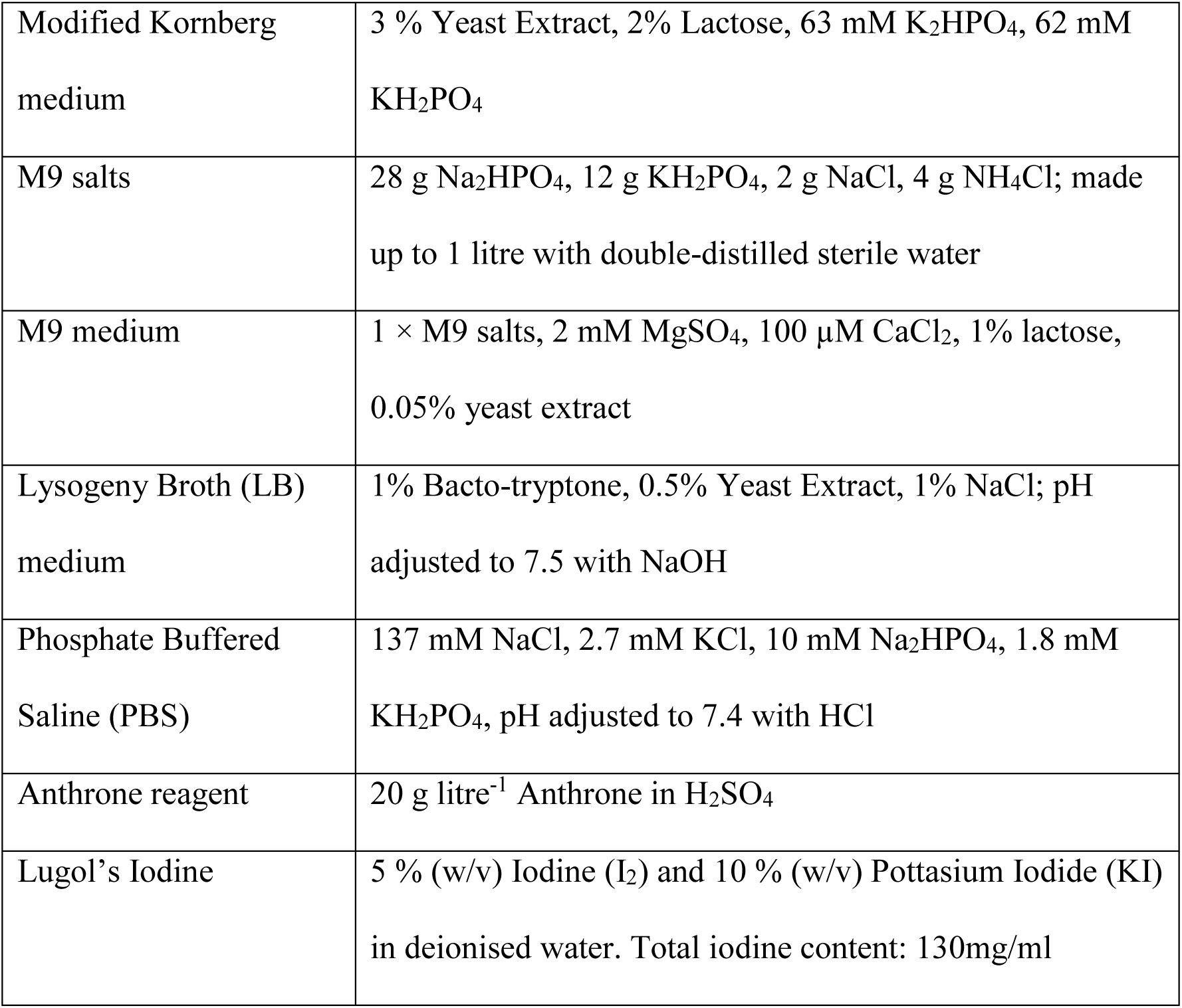
Growth media and assay solutions

|  |  |
| --- | --- |
| Modified Kornberg medium | 3 % Yeast Extract, 2% Lactose, 63 mM K<sub>2</sub>HPO<sub>4</sub>, 62 mM KH<sub>2</sub>PO<sub>4</sub> |
| M9 salts | 28 g Na<sub>2</sub>HPO<sub>4</sub>, 12 g KH<sub>2</sub>PO<sub>4</sub>, 2 g NaCl, 4 g NH<sub>4</sub>Cl; made up to 1 litre with double-distilled sterile water |
| M9 medium | 1 × M9 salts, 2 mM MgSO<sub>4</sub>, 100 µM CaCl<sub>2</sub>, 1% lactose, 0.05% yeast extract |
| Lysogeny Broth (LB) medium | 1% Bacto-tryptone, 0.5% Yeast Extract, 1% NaCl; pH adjusted to 7.5 with NaOH |
| Phosphate Buffered Saline (PBS) | 137 mM NaCl, 2.7 mM KCl, 10 mM Na<sub>2</sub>HPO<sub>4</sub>, 1.8 mM KH<sub>2</sub>PO<sub>4</sub>, pH adjusted to 7.4 with HCl |
| Anthrone reagent | 20 g litre<sup>-1</sup> Anthrone in H<sub>2</sub>SO<sub>4</sub> |
| Lugol's Iodine | 5 % (w/v) Iodine (I<sub>2</sub>) and 10 % (w/v) Pottasium Iodide (KI) in deionised water. Total iodine content: 130mg/ml |

### Growth Curves

Three overnight cultures of LB with chloramphenicol (40 mg/l) were set up for each transformant under investigation, each inoculated with a different colony from a transformation plate. They were incubated overnight at 37 °C and 200 rpm, then OD_λ600nm_ equalised to 3.0 (± 0.09). From each overnight culture, 100 µl was used to inoculate fresh 10 ml cultures of: LB with IPTG (90 mg/l), chloramphenicol (40 mg/l) and lactose (20 g/l) and Kornberg medium with IPTG (90 mg/l), chloramphenicol (40 mg/l) and lactose (20 g/l). The cultures were shaken to mix before 100 µl aliquots were transferred from each fresh culture into the wells of a Costar 3628 flat-bottom 96-well plate in a pattern designed to randomize culture distribution and eliminate edge-effect. The remaining wells surrounding the culture samples were filled with 100 µl of sterile media. In the first instance, 100 µl was transferred from each culture to a 96 well plate twice, using two separate plates that were run simultaneously. In the second instance, only one aliquot was transferred from each culture, so that a single plate was used. The 96-well plates were incubated in a Tecan Sunrise Microplate Absorbance Reader 30041769 at 37 °C and ‘normal’ shaking (4.4 mm, 9.2 Hz). Optical density readings were taken every 15 minutes for 24 hours using the ‘accuracy’ measurement mode. At the same time, the remaining 9.98 ml cultures were incubated at 37 °C and 200 rpm. Optical density readings were taken from these cultures every two hours for the first 14 hours, then at 24 hours. Each optical density reading was obtained by mixing 100 µl of culture with 900 µl of sterile medium in a standard cuvette, then measuring the OD_λ600nm_ against 1 ml sterile medium on a Modulus Single Tube Multimode Reader (Turner Biosystems BS040271) fitted with Absorbance Module E6076.

### Light Microscopy

For each culture, 10 µl was heat-fixed onto a microscope slide. The heat-fixed smear was soaked in 20 µl of 5 % Lugol’s iodine (table 1) for 2 minutes, then rinsed with an excess of 100 % ethanol. Slides were viewed with a Nikon eclipse E200 tabletop microscope under phase contrast at 1000× magnification. Light microscopy images were obtained using a Canon 1XUS9501S digital camera directly through the microscope eyepiece. All images are considered typical of the cultures.

### Transmission Electron Microscopy

For each strain, two 50 ml cultures were grown overnight in LB with IPTG (90 mg/l), chloramphenicol (40 mg/l) and lactose (10 g/l) at 37 °C and 200 rpm. Entire cultures were then spun down at 6000 ×g for 10 minutes. The supernatant was discarded and the pellet washed twice in 25 ml PBS. The cell suspension was then spun down again at 6000 ×g for 10 minutes, the supernatant discarded and the pellet finally re-suspended in 5 ml PBS. From each culture, 1 ml was then transferred to a microcentrifuge tube. These were centrifuged at 6000 ×g for 10 minutes at room temperature and fixed in 3 % glutaraldehyde in buffer C (0.1 M sodium cacodylate buffer, pH 7.3), for 2 hours, then washed in three 10 minute changes of buffer C. Specimens were then post-fixed in 1 % osmium tetroxide in buffer C for 45 minutes, then washed in three 10 minute changes of buffer C. These samples were then dehydrated in 50 %, 70 %, 90 % and 100 % normal grade acetones for 10 minutes each, then for a further two 10-minute changes in analar acetone. Samples were then embedded in araldite resin. Sections, 1 μm thick, were cut on a Reichert OMU4 ultramicrotome, stained with Toluidine Blue, and viewed in a light microscope to select suitable areas for investigation. Ultrathin sections, 60 nm thick, were cut from selected areas, stained in uranyl acetate and lead citrate then viewed in a Philips CM120 Transmission electron microscope. Images were taken on a Gatan Orius CCD camera.

### Iodine Assay

For each culture, 1 ml aliquots were spun down in a tabletop centrifuge at approximately 6000 ×g for 3 minutes at room temperature. Cell pellets were resuspended in 200 µl PBS, to which was added 50 µl of 100 mM CuSO_4_, 50 µl of 6 % v/v H_2_O_2_ and 25 µl of 0.2 % Lugol’s iodine. Suspensions could be transferred to 48-well plates for imaging with a flatbed scanner. Agar plates growing colonies were flooded with 0.2 % Lugol’s iodine solution to test for the presence of starch.

The effect of CuSO_4_ and H_2_O_2_ on the intensity of the stain was investigated by mixing 1 ml starch solution at different concentrations with 25 µl of 0.2 % Lugol’s iodine with or without the prior addition of 50 µl of 100 mM CuSO_4_ and 50 µl of 6 % v/v H_2_O_2_. The optical density of each culture was measured at 620 nm. The soluble starch was from Scientific Laboratory Supplies (CHE3620).

### Anthrone Assays (Individual)

Washed cell pellets were re-suspended in PBS to the volume they had been grown. The optical densities of all cultures were equalised. For each cell suspension, 3 aliquots of 0.33 ml were transferred to microcentrifuge tubes, and to each was added 0.66 ml Anthrone Reagent (table 1). Standard glucose concentration solutions (0 µg ml^-1^, 2 µg ml^-1^, 10 µg ml^-1^, 20 µg ml^-1^, 50 µg ml^-1^ and 100 µg ml^-1^) were also prepared, and for each standard, 3 aliquots of 0.33 ml was mixed with 0.66 ml Anthrone Reagent, in order to provide a standard curve of sugar concentration. The order in which reagent was added to the samples was so arranged that one set of standard glucose solutions was reacted at the start, in the middle, and at the end of the assay process. The order in which reagent was added to each set of culture samples was also alternated. Samples were left on ice for 45 minutes. All samples were then transferred to standard cuvettes and measured at OD_λ620nm_ against the 0 µg ml^-1^ standard sample in a Hitachi Digilab U-1800 spectrophotometer. The glucose equivalent hexose sugar concentration for each sample was estimated against the standard curve of glucose concentrations. Assays therefore measured total hexose sugar content of the cells as a glucose equivalent.

### Starvation Experiment

At the start of the starvation period, cells that had been cultured to maximise their polysaccharide content were recovered by centrifugation, washed and resuspended in M9 with chloramphenicol (40 mg/l) and equalised to an OD_λ600nm_ of 1.5. Each culture was then incubated at 37 °C and 200 rpm, without a carbon source, and sampled at set time points. At each time point, 3 aliquots of 0.33 ml of each culture were transferred to microcentrifuge tubes and assayed as described above. Anthrone assays for the starvation experiment therefore measured the total hexose sugar content of the cultures, to measure how much of the hexose sugar found within the original washed cell pellets at the start of the experiment was metabolised over time.

### SDS Polyacrylamide Gel Electrophoresis

Sonicated culture aliquots containing 30 µg of protein in each case (calculated through Bradford assays) were separated by size on Mini-Protean TGX^TM^ pre-cast SDS polyacrylamide gels (4 – 15%) at 120 V, according to BioRad Laboratories instructions.

### BioBrick™ construction

Plasmid construction conforms to the BioBrick RFC10 method, as first described by Knight (34). The *E. coli* strain JM109, the standard Synthetic Biology plasmid pSB1C3 and the BioBrick BBa_K523005 (P_LAC_-RBS-LacZ-RBS) were used for all cloning procedures and the propagation of plasmid DNA. For *glgC* and *glgC16*, BioBrick parts BBa_K118015 and BBa_K118016 (respectively) were used, giving the genes with *EcoRI* sites removed and the G336D substitution in the case of *glgC16*. Table 2 shows the primers used to amplify *glgB* and *glgC* from the *E. coli* chromosome, as well as those used to remove the *EcoRI* sites (glgCm1 and glgCm2). All restriction enzymes were purchased from New England Biolabs (NEB) and used according to the manufacturer’s instructions.

**Table 2:**
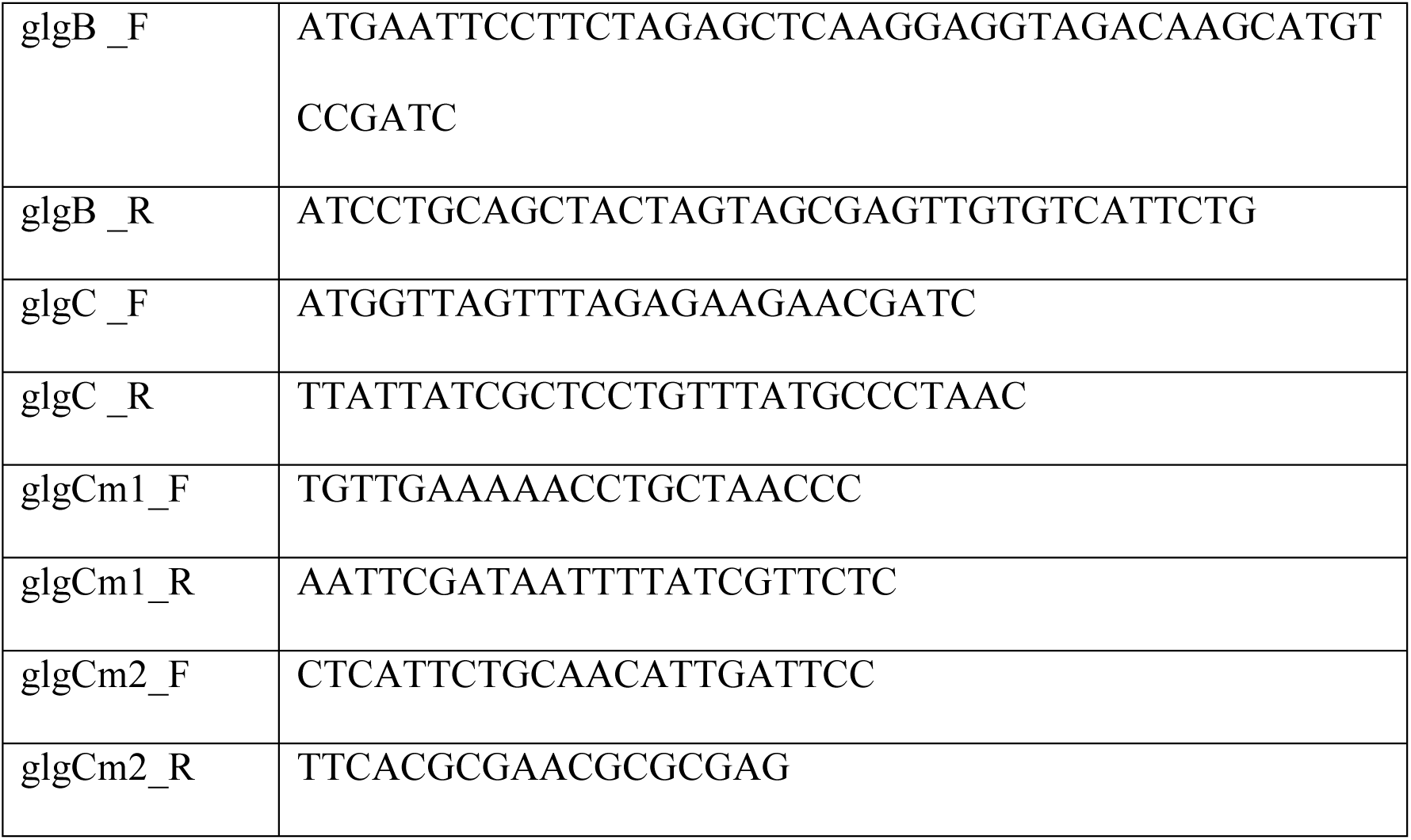
Primers

**Table 3:** Plasmids

| Plasmid | Description | Source |
| --- | --- | --- |
| pJW-lacZ | pSB1C3 + lacZ-α | This work |
| pJW-glgC | pSB1C3 + lacZ-α + *glgC* | This work |
| pJW-glgC16 | pSB1C3 + lacZ-α + *glgC16* | This work |
| pJW-glgB | pSB1C3 + lacZ-α + *glgB* | This work |
| pJW-glgCB | pSB1C3 + lacZ-α + *glgC* + *glgB* | This work |

## Acknowledgments

This work was supported by the Biotechnology and Biological Sciences Research Council, grant number BB/F017073/1.

## References

1. Conca J. 2014. It’s final - corn ethanol is of no use. Forbes.com. https://www.forbes.com/sites/jamesconca/2014/04/20/its-final-corn-ethanol-is-of-no-use/#.

2. Belkin B. 2008. Food inflation, riots spark worries for world leaders. WSJ. https://www.wsj.com/articles/SB120813134819111573

3. Sachs J. 2008. Surging food prices and global stability. Scientific American 298:40–40.

4. Fargione J, Hill J, Tilman D, Polasky S, Hawthorne P. 2008. Land clearing and the biofuel carbon debt. Science 319:1235–1238.

5. Searchinger T, Heimlich R, Houghton R, Dong F, Elobeid A, Fabiosa J, Tokgoz S, Hayes D, Yu T. 2008. Use of U.S. croplands for biofuels increases greenhouse gases through emissions from land-use change. Science 319:1238–1240.

6. Lynd L, Zyl W, McBride J, Laser M. 2005. Consolidated bioprocessing of cellulosic biomass: an update. Current Opinion in Biotechnology 16:577–583.

7. Kumar R, Singh S, Singh O. 2008. Bioconversion of lignocellulosic biomass: biochemical and molecular perspectives. Journal of Industrial Microbiology & Biotechnology 35:377–391.

8. Wackett L. 2008. Biomass to fuels via microbial transformations. Current Opinion in Chemical Biology 12:187–193.

9. Yuan J, Tiller K, Al-Ahmad H, Stewart N, Stewart C. 2008. Plants to power: bioenergy to fuel the future. Trends in Plant Science 13:421–429.

10. French C. 2009. Synthetic biology and biomass conversion: a match made in heaven?. Journal of The Royal Society Interface 6.

11. Taha M, Foda M, Shahsavari E, Aburto-Medina A, Adetutu E, Ball A. 2016. Commercial feasibility of lignocellulose biodegradation: possibilities and challenges. Current Opinion in Biotechnology 38:190–197.

12. Falcón L, Lindvall S, Bauer K, Bergman B, Carpenter E. 2004. Ultrastructure of unicellular N2-fixing cyanobacteria from the tropical North Atlantic and subtropical North Pacific oceans. Journal of Phycology 40:1074–1078.

13. Deschamps P, Colleoni C, Nakamura Y, Suzuki E, Putaux J, Buleon A, Haebel S, Ritte G, Steup M, Falcon L, Moreira D, Loffelhardt W, Raj J, Plancke C, d’Hulst C, Dauvillee D, Ball S. 2008. Metabolic symbiosis and the birth of the plant kingdom. Molecular Biology and Evolution 25:795–795.

14. Ball S, Morell M. 2003. From bacterial glycogen to starch: understanding the biogenesis of the plant starch granule. Annual Review of Plant Biology 54:207–233.

15. D’Hulst C, Mérida Á. 2010. The priming of storage glucan synthesis from bacteria to plants: current knowledge and new developments. New Phytologist 188:13–21.

16. Smith A. 1999. Making starch. Current Opinion in Plant Biology 2:223–229.

17. Smith A. 2001. The biosynthesis of starch granules. Biomacromolecules 2:335–341.

18. Zeeman S, Kossmann J, Smith A. 2010. Starch: its metabolism, evolution, and biotechnological modification in plants. Annual Review of Plant Biology 61:209–234.

19. Tetlow I. 2006. Understanding storage starch biosynthesis in plants: a means to quality improvement. Canadian Journal of Botany 84:1167–1185.

20. Saibene D, Zobel H, Thompson D, Seetharaman K. 2008. Iodine-binding in granular starch: different effects of moisture content for corn and potato starch. Starch - Stärke 60:165–173.

21. Yu X. 1996. The complex of amylose and iodine. Carbohydrate Research 292:129–141.

22. John M, Schmidt J, Kneifel H. 1983. Iodine—maltosaccharide complexes: relation between chain-length and colour. Carbohydrate Research 119:254–257.

23. McGrance S, Cornell H, Rix C. 1998. A simple and rapid colorimetric method for the determination of amylose in starch products. Starch - Stärke 50:158–163.

24. Moulay S. 2013. Molecular iodine/polymer complexes. Journal of Polymer Engineering 33.

25. Meléndez R, Meléndez-Hevia E, Mas F, Mach J, Cascante M. 1998. Physical constraints in the synthesis of glycogen that influence its structural homogeneity: a two-dimensional approach. Biophysical Journal 75:106–114.

26. Ball S, Colleoni C, Cenci U, Raj J, Tirtiaux C. 2011. The evolution of glycogen and starch metabolism in eukaryotes gives molecular clues to understand the establishment of plastid endosymbiosis. Journal of Experimental Botany 62:1775–1801.

27. Wilson W, Roach P, Montero M, Baroja-Fernández E, Muñoz F, Eydallin G, Viale A, Pozueta-Romero J. 2010. Regulation of glycogen metabolism in yeast and bacteria. FEMS Microbiology Reviews 34:952–985.

28. Govons S, Gentner N, Greenberg E, Preiss J. 1973. Biosynthesis of bacterial glycogen XI. Kinetic characterization of an altered adenosine diphosphate-glucose synthase from a ‘glycogen-excess’ mutant of *Escherichia coli* B. Journal of Biological Chemistry 248:1731–1740.

29. Manonmani H, Kunhi A. 1999. Interference of thiol-compounds with dextrinizing activity assay of α-amylase by starch-iodine colour reaction: modification of the method to eliminate this interference. World Journal of Microbiology and Biotechnology 15:485–487.

30. Zeeman S, Delatte T, Messerli G, Umhang M, Stettler M, Mettler T, Streb S, Reinhold H, Kötting O. 2007. Starch breakdown: recent discoveries suggest distinct pathways and novel mechanisms. Functional Plant Biology 34:465.

31. Mahlow S, Orzechowski S, Fettke J. 2016. Starch phosphorylation: insights and perspectives. Cellular and Molecular Life Sciences 73:2753–2764.

32. Chung C, Niemela S, Miller R. 1989. One-step preparation of competent *Escherichia coli*: transformation and storage of bacterial cells in the same solution. Proceedings of the National Academy of Sciences 86:2172–2175.

33. Sundberg M, Pfister B, Fulton D, Bischof S, Delatte T, Eicke S, Stettler M, Smith S, Streb S, Zeeman S. 2013. The heteromultimeric debranching enzyme involved in starch synthesis in Arabidopsis requires both isoamylase1 and isoamylase2 subunits for complex stability and activity. PLoS ONE 8:e75223.

34. Knight, T., 2003. Idempotent vector design for standard assembly of biobricks. Massachusetts inst. of tech Cambridge artificial intelligence lab.

